# Post-movement beta rebound reflects strategic re-aiming during motor adaptation, but not re-aiming accuracy

**DOI:** 10.1101/2025.01.17.633560

**Authors:** Matthias Will, Betina Korka, Max-Philipp Stenner

## Abstract

Motor adaptation results from several interacting learning mechanisms, including learning via cognitive strategies and implicit adaptation. While strategy-based and implicit learning can be dissociated at a behavioural level, their underlying systems-level physiology is poorly understood. A neural signal that undergoes pronounced changes during motor adaptation is the post-movement beta-rebound (PMBR). However, it is unclear how these changes relate to the specific learning mechanisms that contribute to motor adaptation. We measured electroencephalography (EEG) while healthy participants (N=27) performed reaching movements towards a target. In most trials, a cursor showed the veridical position of the unseen hand, however, for some reaches, the direction of the cursor was rotated relative to the position of the hand. Participants were informed that, once a rotation occurred, it could persist for a single trial (1x condition), or for two consecutive trial (2x condition). In the 2x condition, participants could therefore redirect the rotated cursor through the target in the second rotated trial by re-aiming, while they had to continue aiming at the target in the 1x condition. We observed a stronger decrease of PMBR following the first rotated reach in the 2x condition, compared to the 1x condition, despite similar kinematics. This corroborates our previous results that PMBR reflects strategic re-aiming (Korka *et al*., 2023). However, when we collapsed data from the two studies (total N=53), we found that the degree to which the PMBR decreases does not predict re-aiming accuracy. We discuss the role of PMBR in motor adaptation, including implications for clinical disorders.

## INTRODUCTION

In order to achieve our goals, we continuously adapt our movements to a changing environment. Motor adaptation arises from several learning mechanisms, including implicit adaptation and cognitive strategies. Implicit motor adaptation is a highly automatic learning process that is thought to reduce any error in predicting the sensory consequences of movement via adjustments to an internal model (Wolpert, Ghahramani and Jordan, 1995; Mazzoni and Krakauer, 2006; Tseng *et al*., 2007). This sensory prediction error can be introduced, for example, by a visuomotor perturbation. Given its high degree of automaticity, a hallmark of implicit adaptation are aftereffects, i.e., persistent adaptation for a short period after a perturbation has been removed. Strategy-based adaptation, on the other hand, is a deliberate, cognitively more effortful process that aims to reduce target error (i.e., errors in achieving the task goal, for example, hitting a movement target) through the selection of alternative actions, such as by re-aiming (Taylor, Krakauer and Ivry, 2014; McDougle and Taylor, 2019). Several experimental manipulations can dissociate implicit adaptation and strategy-based learning at a behavioral level (Haith, Huberdeau and Krakauer, 2015; Morehead *et al*., 2016; Hadjiosif and Krakauer, 2021; Maresch, Werner and Donchin, 2021). However, the neural circuitry that gives rise to different motor adaptation mechanisms is not yet well understood.

Motor adaptation is accompanied by typical modulations of neuronal population signals that can be captured using EEG (Reuter, Booms and Leow, 2022). However, the precise information content of these signal modulations with respect to the mental operations involved in motor adaptation is still unclear. Among these signals, movement-related modulations of oscillations in the beta band (13-30 Hz) have attracted considerable attention (Rustighi, Dohnal and Mace, 2010; Spapé and Serrien, 2010; Rilk *et al*., 2011; Savoie *et al*., 2018; Alayrangues *et al*., 2019; Jahani *et al*., 2020). Beta power displays prominent modulations around the time of movement (Kilavik *et al*., 2013). It decreases before and during movement, and rebounds shortly after movement offset (post-movement beta rebound, PMBR). When visual feedback for movement is perturbed, PMBR is reduced. However, it is debated how exactly this modulation relates to motor adaptation. For example, Tan et al. (Tan, Jenkinson and Brown, 2014; Tan, Wade and Brown, 2016) have proposed that the decrease in PMBR when a perturbation is first introduced mirrors a decrease in confidence in internal models, driven by sensory prediction error. Tan *et al*. thus suggested a link between PMBR and a key mechanism thought to underlie implicit motor adaptation. Torrecillos *et al*. (Torrecillos *et al*., 2015), on the other hand, observed a decrease in PMBR even in response to a target jump, an experimental manipulation that does not produce any implicit motor adaptation. They suggested, therefore, a more general role of PMBR in the processing of salient events.

It is possible that PMBR is modulated by multiple aspects of motor adaptation (Kilavik *et al*., 2013), and that the extent of modulation depends on the types of learning involved in the task at hand. The studies by Tan *et al*. (2014, 2016) employed large visuomotor perturbations that remained constant for many consecutive trials, which typically emphasizes strategy-based learning (Morehead *et al*., 2015). The design of Torrecillos *et al*. (2015), on the other hand, prevented strategy-based learning because perturbations were always switched off after a single trial. In a direct comparison of motor adaptation with vs. without strategic re-aiming, we have recently shown that the decrease in PMBR in response to a newly introduced visuomotor perturbation depends on whether, or not, subjects employ a re-aiming strategy (Korka *et al*., 2023). Specifically, the decrease in PMBR is more pronounced when subjects re-aim, compared to purely implicit adaptation.

Here, we replicate this finding, using a paradigm that further de-confounds re-aiming and the type of perturbation. In our previous study, we had compared adaptation to a clamped rotation (or error-clamp), which renders re-aiming futile, with adaptation to a visuomotor rotation, which allows for re-aiming. Error-clamping entails visual feedback that moves always in the same direction relative to the target, irrespective of the direction of hand movement. While this can effectively prevent re-aiming, error-clamping can also lead to a distortion in the mapping of hand trajectories to visual feedback, for example, when the hand follows a curved path, while the clamped visual feedback remains always straight. To avoid any difference in the relation between movement and feedback between conditions, the present study employed the same type of perturbation, i.e., a visuomotor rotation, across conditions, and prevented re-aiming purely through prior knowledge of the sequence of perturbations.

The second goal of the present study was to reveal whether inter-individual differences in the modulation of PMBR predict inter-individual differences in strategic re-aiming. Individuals with Parkinsońs disease and cerebellar degeneration have greater difficulties discovering a re-aiming strategy than healthy controls (Butcher *et al*., 2017; Wong *et al*., 2019; Tsay, Najafi, *et al*., 2022), and also display a reduced PMBR (after unperturbed hand movements; Pfurtscheller *et al*., 1998; Tamás *et al*., 2003; Aoh *et al*., 2019; Visani *et al*., 2020). This raises the interesting possibility that PMBR, or its modulation by error or sensorimotor perturbation, may reflect a person’s capacity for strategy-based learning. To explore this possibility in a healthy cohort, we took advantage of the highly similar design of the re-aiming condition in the present study, and our previous study (Korka *et al*., 2023). This high similarity allowed us to collapse data across our previous study, and the present study, and relate PMBR, and its modulation by a visuomotor rotation, to re-aiming accuracy in a relatively large cohort of 53 subjects.

## MATERIALS AND METHODS

### Participants

30 healthy right-handed volunteers completed the experiment. Handedness was determined via the Edinburgh Handedness Inventory (Oldfield, 1971). Three participants did not follow re-aiming instructions, evident in a high number of rejected trials (see ‘Analysis of kinematic data’ for rejection criteria), and were thus excluded. The final sample size of 27 participants included 17 females and 10 males, and was, on average, 28.4 years old. The sample size was chosen to match the sample size in our previous study (Korka *et al*., 2023, N=26, 14 female, 23.8 years mean age). Participants gave written informed consent, and the study was approved by the ethics committee of the Medical Faculty of the University Magdeburg, and conducted in accordance with the Declaration of Helsinki. Participants received monetary compensation.

### Apparatus

Participants were sitting in a quiet, dimly lit, electrically shielded chamber (Industrial Acoustics Company). Both in this study, and our previous study (Korka *et al*., 2023), participants moved a stylus held in their right hand across a graphics tablet (Wacom Intuos Pro Large, Kazo, Japan; sampling rate of 200 Hz, active area of 311 × 216 mm). The stylus tip was continuously in contact with the tablet, which recorded its position. An LCD screen (refresh rate 60 Hz) mounted horizontally above the tablet occluded vision of the hand, and provided visual feedback (see **Fig. 1A**).

**Fig. 1:**
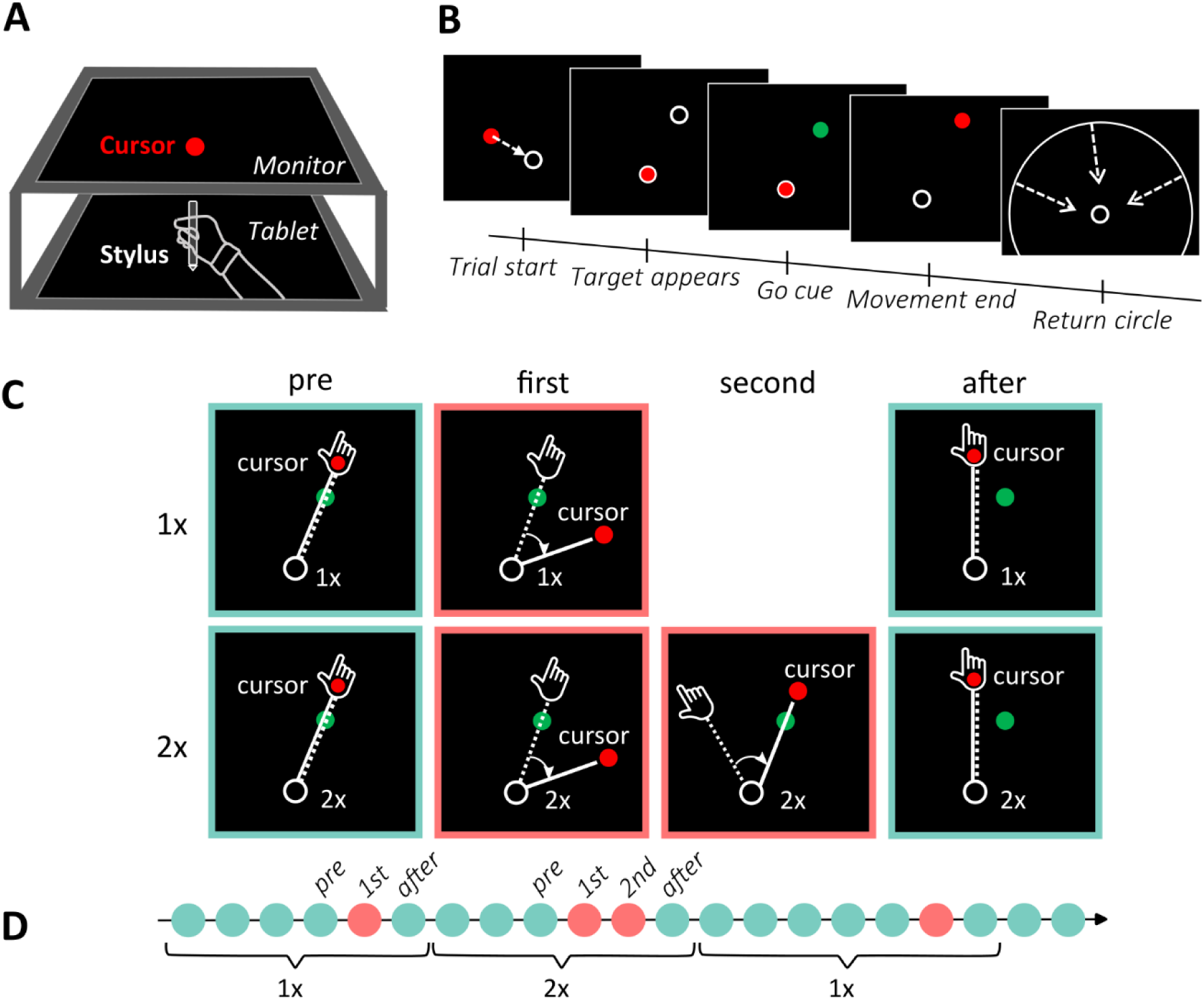
Set-up and task. (**A**) The experimental set-up consisted of a tablet, on which participants performed center-out reaching movements holding a stylus, and a monitor mounted above the tablet, on which online visual feedback was displayed. (**B**) Participants moved the cursor (red dot) to the home position (white circle) at the beginning of each trial. After holding the stylus in place for 2000-2500 ms, the target appeared (white circle). Once the target turned green (1500 ms after target onset), participants could initiate a reaching movement to slice the cursor through the target. After movement offset, participants held their hand in the endpoint position for a short duration (1600-2100 ms). A white circle, whose radius corresponded to the radial distance of the hand from the home position, guided the return movement. (**C**) Cursor and hand position were identical on most trials (cyan frame). On some trials, the cursor trajectory was rotated relative to hand movement (red frame). In each cycle, there could be a single trial with a rotation (1x condition, upper row), or two consecutive trials with identical rotation (2x condition, lower row). In both conditions, the first trial with a rotation was not predictable, so that participants could not anticipate the rotation, and the cursor missed the target on the first rotated trial. Participants were instructed to compensate for the rotation in the second trial with a rotation in the 2x condition by re-aiming their reach in the opposite direction of the rotation. Following the second rotated trial (2x condition) or single rotated trial (1x condition), cursor and hand position were again aligned, and participants again aimed directly at the target (after-trial). We expected that participants would implicitly adapt their movements to the visuomotor rotation in both conditions and consequently show an after-effect in the after trial, evident in slightly missing the target in the direction opposite to the rotation. (**D**) Conditions alternated after every cycle.

### Task and procedure

Participants performed center-out reaching movements with their dominant right hand in a horizontal plane to make a cursor on the screen “slice” through a visual target. Stimuli and events in individual trials were identical to our previous study (Korka *et al*., 2023). Trials started with the hand holding the stylus still in the home position (white circle, outer diameter of 8.5 mm, see **Fig. 1B**). After a variable time interval between 2000 and 2500 ms (uniform random distribution), the target (white circle, outer diameter of 6 mm) appeared 2 cm away from the home position at an angle of 45° clockwise relative to the “straight ahead” direction. Participants were instructed to initiate the movement once the target turned green (1500 ms after target onset), with no time limit for movement initiation after that. Movement initiation was defined as the time point when the tip of the stylus left the home position. Participants had to cross the target radius within a time window of 50 to 160 ms after movement initiation. If they crossed the target distance earlier than 50 ms, or later than 160 ms, or if they initiated their movement before the target turned green, they saw a corresponding error message on the screen (“TOO FAST”, “TOO SLOW”, or “TOO EARLY”, respectively). Throughout the movement, participants saw a cursor (red dot, diameter 5 mm) as visual feedback of their movement. After stopping their movement behind the target (with movement offset defined as the point in time when the cursor did not change position for at least two vertical frame refreshes, i.e., 33.3 ms), participants had to hold the stylus in place (in the endpoint position) for at least 1600 to 2100 ms (uniform random distribution). If participants moved their hand more than 5 mm away from the endpoint position during this period, they received an error message (“HOLD STILL”). They then moved their hand back to the home position, guided by a circle whose radius corresponded to the distance of the hand from the home position. This prevented feedback about the angular position of their hand while providing information about the radial distance from the home position. The trial ended when the hand reached the home position (see **Fig. 1B**).

The experiment consisted of 16 blocks of 72 trials each over the course of two days of testing (8 blocks per day). Trials in each block were organized in 12 cycles of different lengths (at least 4 trials/cycle). In most trials (54 per block), cursor and hand position were aligned (unrotated trials). In the other, pseudo-randomly interspersed trials, the movement of the cursor was rotated relative to the movement of the hand (rotated trials, 18 per block). There were two conditions: each cycle contained either a single trial with a rotation (1x condition), or two consecutive trials with a rotation (2x condition; see **Fig. 1C**). Throughout each cycle, participants saw an indicator on the screen showing the current condition (“1x” or “2x”). Thus, they knew beforehand whether any rotation they encountered would be repeated once in the subsequent trial (2x), or not (1x). To minimize the possibility that participants used a single, fixed strategy throughout the experiment, rotations differed in magnitude (30°, 37.5°, or 45°) and direction (clockwise, CW, or counterclockwise, CCW) across cycles. Each of the three rotation magnitudes and two rotation directions occurred in an equal number of cycles per block, and equally often in cycles of each condition. Importantly, in the 2x condition, the two consecutive, rotated trials had the same rotation magnitude and direction. This allowed participants to re-aim their reach in the second rotated trial, based on the rotation experienced in the first rotated trial, so that the cursor would slice through the target. Due to variable cycle length, participants could not form any reliable prediction about when a rotation was going to occur for the first time in a cycle. We therefore expected participants to follow the default instruction to aim their movement directly at the target, except in the second rotated trial in the 2x condition. After the last rotated trial in each cycle, there was a further unrotated trial, in which participants were instructed to aim directly at the target again. Following that trial, the indicator on the screen changed (from “1x” to “2x”, or back), prompting the change in condition (see **Fig. 1D**).

Similarly to the present study, participants in our previous study (Korka *et al*., 2023) had to ignore or compensate for a visuomotor perturbation. There were two conditions with different types of visuomotor perturbations. In contrast to the present study, the *Ignore* condition used movement direction-invariant (or clamped) visual feedback, rendering any re-aiming useless (Morehead *et al*., 2016). The other condition (*Compensate*) was virtually identical to the 2x condition in the present study, with the only exception that participants completed entire blocks of trials of the *Compensate* condition, rather than shorter cycles of trials (for details see Korka *et al*., 2023). For the median split analysis in the present manuscript, we only include data from the *Compensate* condition.

### Analysis of kinematic data

Data analysis (behavioral and EEG) for the present study was similar to our previous study (Korka *et al*., 2023). The primary kinematic variable of interest in this study was movement direction relative to the target at the point of maximum velocity. Movement direction for rotated trials and corresponding after-effect trials was defined as positive for the direction opposite to the rotation. We also computed response time, movement time, movement extent, and movement curvature. Response time was defined as the duration between the Go signal (the target turning green) and movement initiation. Note that participants could plan a movement earlier than the Go signal because the target was displayed before, and target location remained constant throughout the experiment. Movement curvature was computed as the maximum perpendicular distance between a movement trajectory and a straight line from the home position to the movement endpoint, relative to movement extent (Atkeson and Hollerbach, 1985).

We excluded all trials in which participants received an error message (“TOO SLOW”, “TOO FAST”, “TOO EARLY”, and “HOLD STILL”), as well as trials that followed trials with one of the former two error messages. Finally, we excluded cycles in which participants did not follow the aiming instructions, i.e., cycles in the 2x condition in which movement angle in the second rotated trial was <15° away from the target, and cycles in the 1x condition in which movement angle in the trial following the single rotated trial was >15° away from the target, i.e., we used the same cutoff of movement direction as in our previous study (Korka *et al*., 2023).

We report results for the pre-trial (trial before the first rotation in each cycle), first rotated trial, second rotated trial, and after-trial (first unrotated trial after the rotation in each cycle, see **Fig. 1D**).

### EEG data recording

In each session, EEG was recorded using 35 Ag-AgCl passive electrodes connected to a BrainAmp amplifier, with data acquisition handled by the Vision Recorder software (Brain Products). The sampling rate was set at 500 Hz. Two electrodes were positioned on the mastoids, with the right mastoid serving as the online reference. The ground electrode was placed at the Fpz location. Three electrodes were used for recording EOG activity: two were placed on the outer canthi of the left and right eyes, and one below the left eye. The remaining 29 electrodes (Fp1/2, F3/4, F7/8, FC1/2, C3/4, CP1/2, T7/8, P3/4, P7/8, PO3/4, PO7/8, O9/10, Fz, Cz, Pz, Oz, Iz) were mounted in an elastic cap (EasyCap) according to the extended international 10-20 system. Before beginning data recording, we ensured that all electrode impedances were below 10 kΩ. The raw EEG data are available online under https://doi.org/10.5281/zenodo.14673450 and https://doi.org/10.5281/zenodo.14674428 (together with the kinematic data https://doi.org/10.5281/zenodo.14671785).

### Analysis of EEG data

The signal of interest for EEG analysis was the PMBR. EEG pre-processing was performed in MATLAB R2020b (Mathworks) using the FieldTrip toolbox (Oostenveld *et al*., 2011). First, the continuous raw data was filtered using a 100 Hz low-pass and a 0.1 Hz high-pass windowed sinc finite impulse response (FIR) filter (Hamming window, filter order 66-low-pass and 8250-high-pass). Next, the continuous data set was epoched, with epochs starting 2 s before target onset and ending 2 s after beginning of the return movement. We removed epochs with exceptionally high amplitude fluctuations. Specifically, for each epoch we computed the ‘robust standard deviation’ (i.e., 0.7413 times the inter-quartile-range; Bigdely-Shamlo *et al*., 2015) of EEG amplitudes across time for each channel, and then calculated the median robust standard deviation across all channels. Epochs were rejected using a thresholding approach based on the ‘robust z score’ (Bigdely-Shamlo *et al*., 2015) of the obtained epoch distribution (robust z>6). Thresholds were chosen to strike a balance between maintaining enough data and rejecting clear outliers, which was confirmed by visual inspection. We removed 11.1 trials on average (range 0 to 37). Similarly, we computed the robust standard deviation after concatenating all trials, for each channel separately. Noisy channels were identified using the robust z score of the channel distribution (robust z>3) and removed. On average we removed 1.1 channels (range: 0 to 4). Rejected channels were replaced with a weighted average of all neighbors. Following this, we ran an ICA on a copy of the data. This copy was processed in a similar way as described above with the difference that the applied high-pass filter used a higher cut-off frequency of 1 Hz (windowed sinc FIR filter, filter order 826, same low-pass filter as described before) to optimize identification of eye movement artifacts via ICA. We also used a more rigorous trial rejection threshold for this copy of the dataset (robust z>3) to avoid any bias to the ICA due to artifactual trials. Components related to eye movements were identified via visual inspection and the obtained mixing and unmixing matrices were applied to the original dataset, removing the identified eye movement components. As the final step in the pre-processing pipeline, we re-referenced to a common average across all electrodes (not including EOG).

We performed a time-frequency analysis (*ft_freqanalysis* function in FieldTrip with ‘mtmconvol’ method and a 400 ms Hanning taper) from which we obtained spectral power for frequencies from 2.5 Hz to 40 Hz (in steps of 2.5 Hz) and time points from-3200 ms to 2200 ms relative to movement offset (in steps of 50 ms). A shared baseline for the 1x and 2x condition was defined as the spectral power during the inter-trial interval (period of-1800 ms to-200 ms relative to target onset, spectral power averaged over time) before unrotated trials (except pre-and after-trials) from both conditions. Spectral power was averaged across trials, separately for each condition, i.e. pre-trials, first and second rotation, and after-trials, each for the 1x condition and 2x condition. Next, we identified channels of interest for PMBR analysis. Importantly, channel identification was performed only on unrotated trials (except pre-and after-trials), i.e., trials that are not included in the statistical PMBR analysis, which is therefore unbiased by this selection step. We defined channels of interest based on visual inspection by selecting channels that showed a prominent PMBR, evident as an increase in spectral power in the beta-frequency range after movement offset relative to baseline (channel labels in black font in **Fig. 2A**). Subsequently, we refined the range for the time and frequency of interest by inspecting spectral power averaged over the selected channels and identified a time-frequency window of interest that ranged from 15 Hz to 30 Hz, and from 200 ms to 1200 ms after movement offset (see **Fig. 2B**). We verified this selection of channels and time-frequency window of interest by comparison with a positive cluster obtained via FieldTrip’s cluster-based permutation test, which compared beta-power in a broader post-movement time-frequency window (12.5 Hz to 40 Hz, 100 ms to 1800 ms) and across all channels to baseline power (as defined above). This test revealed a time-and frequency-range that was similar to the selection by visual inspection, and a slightly larger number of channels that displayed an increase in post-movement beta power, which included more frontal and parietal channels in addition to the channels of interest identified via visual inspection (channel labels in grey and black font, respectively, in **Fig. 2A**). Subsequent PMBR analyses were based on the selection of channels of interest by visual inspection, which matched better with the topography found in our previous study (Korka *et al*., 2023). However, results are qualitatively identical when using the larger selection of channels of interest. Finally, we obtained a single value as a measure for the PMBR for each trial type (pre, first, second, and after) and condition (1x and 2x) by averaging the change in spectral power relative to baseline across the channels and time-frequency window of interest.

**Fig. 2:**
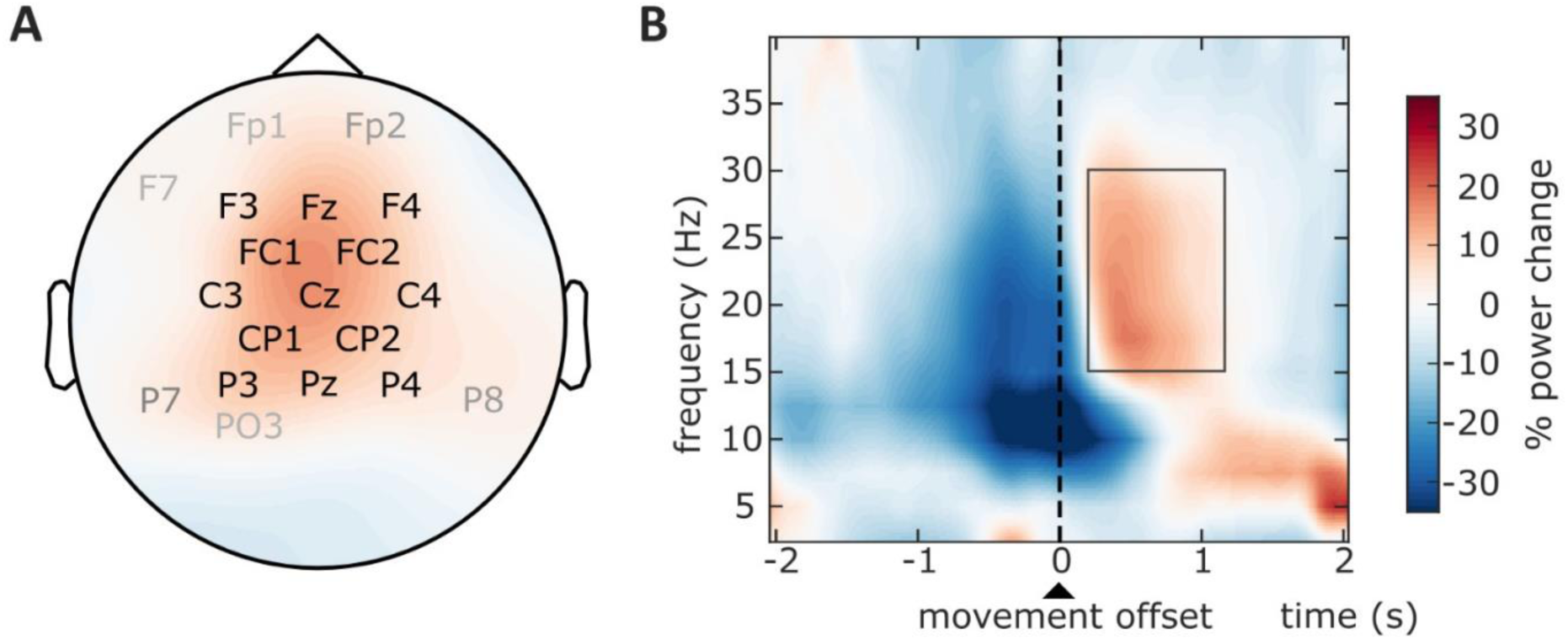
Channels and time-frequency window of interest. (**A**) Channels identified by visual inspection (black labels) and permutation-test (grey labels). The color codes for power change, in the time-frequency window of interest from 200 ms to 1200 ms and 15 Hz to 30 Hz, relative to baseline (1800 ms to 200 ms before target onset, same frequency window) (**B**) Power change relative to baseline, averaged across the channels of interest shown in black in panel A. The black rectangle indicates the time-frequency window of interest.

### Median split

After we confirmed that PMBR was modulated by use of an aiming strategy, we asked if individual differences in the decrease in PMBR from pre-to first rotated trial could predict re-aiming accuracy. To this end, we performed a median split between participants based on the average unsigned target error for the second rotated trial, as a measure for re-aiming performance. For this analysis we included data from a re-aiming condition from our previous study, which was virtually identical to the 2x condition of the present study. The only difference between both datasets is that the previous study used a blocked design, while conditions were interleaved in the present study. Participants of both experiments were balanced across splits. Thus, any differences in PMBR across experiments could not explain potential differences across splits. A total of 53 individuals contributed data for the median-split analysis.

### Statistics

We report mean values and standard deviation or median, and inter-quartile range (IQR), when a Shapiro-Wilk test indicated a violation of the assumption of normality. Movement kinematics between conditions were compared using paired t-tests. When the assumption of normality was violated, we used a corresponding nonparametric test. Bayesian t-tests were used to test for evidence in favor of the null hypothesis. Repeated-measures analyses of variance (ANOVA) were used to test for differences in movement direction and differences in PMBR between conditions and trial types. For the median-split analysis, we employed a mixed-ANOVA. Statistics were computed in MATLAB R2020b and JASP 0.14.0.0 (JASP Team, 2020).

## RESULTS

### PMBR is modulated by strategy-based motor adaptation

Our primary hypothesis was that the decrease in PMBR during visuomotor adaptation would be modulated by the behavioral relevance of movement errors, that is, the error between the rotated cursor and target (target error). We expected a more prominent reduction in PMBR on the first rotated trial, relative to the pre-trial, when the target error was relevant for the next movement, i.e., when it was informative for devising a re-aiming strategy (2x condition), compared to when it was not relevant for the next movement, i.e., when there was no deliberate change in behavior following the first rotated trial (1x condition).

We first verified that there were no condition differences in kinematics in the first rotated trial that could confound the EEG analysis. There were no significant differences in response time (1x: 0.55 ± 0.15 s; 2x: 0.56 ± 0.15 s; mean ± SD), movement time (1x: 0.31 ± 0.04 s, 2x: 0.31 ± 0.04 s), movement extent (1x: 4.00 ± 0.98 cm, 2x: 4.00 ± 1.01 cm), maximum movement velocity (1x: 24.72 ± 4.52 cm/s, 2x: 24.67 ± 4.30 cm/s) and movement curvature (linearity index, 1x: 0.09 ± 0.03, 2x: 0.09 ± 0.03) between the 1x and 2x condition (all t(26)<0.89, all p>0.38). Bayesian paired t-tests provided moderate evidence that kinematic parameters were indeed similar between conditions (all BF_10_<0.3).

Next, we examined movement direction across different trial types and conditions. Across the 1x and 2x condition, participants were similarly accurate in slicing through the target during the pre-trial (1x:-0.21 ± 0.66°, 2x: 0.14 ± 0.62°; W=120, p=0.10, BF_10_=1.02) and first rotated trial (1x:-0.11 ± 0.44°, 2x: 0.12 ± 0.58°; W=143, p=0.28, BF_10_=0.53; **Fig. 3A**). In the 2x condition, participants compensated for the rotation in the second rotated trial by aiming in the opposite direction, to a degree that was sensitive to the magnitude of rotation (**Fig. 3B**). A rANOVA revealed a main effect of Rotation Magnitude (30°, 37.5°, 45°) on movement direction (31.66 ± 4.94°, 34.87 ± 4.83°, 40.40 ± 7.00°, respectively; F(2,52)=26.3, p<0.001, η^2^=0.50). Post-hoc tests showed that movement direction differed between all rotation magnitudes (30° vs 37.5°: t(26)=2.64, p=0.011, d=0.57; 30° vs 45°: t(26)=7.17, p<0.001, d=1.54; 37.5° vs 45°: t(26)=4.53, p<0.001, d=0.97; all Holm corrected).

**Fig. 3:**
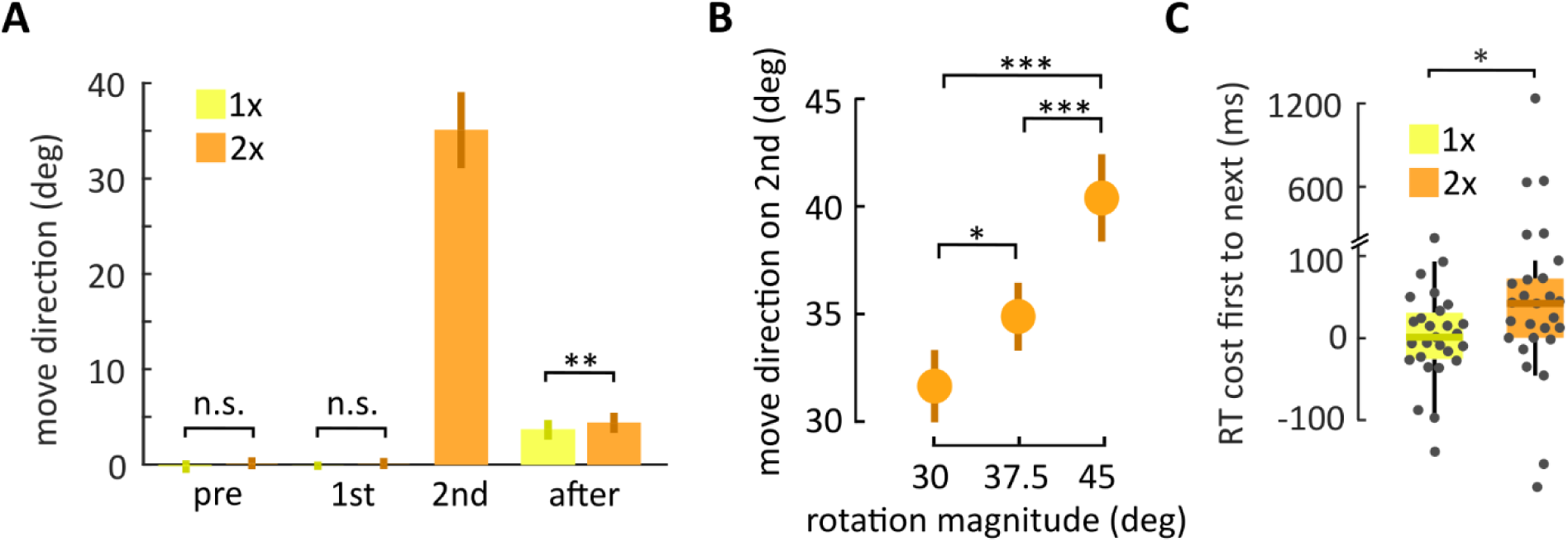
Behavioral results. (**A**) Mean movement direction (bars, vertical bold lines indicate standard deviation) across different trials. The y-axis shows movement direction relative to the target. Positive values on the y-axis indicate movement angles opposite to the rotation that participants experienced in the first rotated trial (and, in the 2x condition, in the second rotated trial). Participants moved their hand to the target during the pre-trial and first rotated trial in the 1x (yellow) and 2x condition (orange). When a second, identical rotation followed the first rotation, i.e., in the 2x condition, participants compensated for the angular deviation by aiming in the opposite direction. On the trial after the rotation, participants were biased in the direction opposite to the previously experienced rotation due to implicit adaptation. (**B**) Aiming direction in the second rotated trial was selected according to the rotation magnitude (dots indicate mean, vertical bold lines indicate standard deviation). (**C**) Re-aiming was associated with a higher response time cost compared to maintaining aim directly at the target (boxplots, dots represent individual participants).

A 2×2 rANOVA indicated a significant difference in response times (RT) for the factor Condition (1x and 2x; F(1,26)=4.65, p=0.040, η^2^=0.05) and a trend for the factor Trial Type (first rotated trial and next trial, i.e., after-trial in 1x condition and second rotated trial in 2x condition; F(1,23)=3.96, p=0.056), as well as an interaction between both factors (F(1,26)=4.80, p=0.038, η^2^=0.04; **Fig. 3C**). In agreement with increased cognitive load during re-aiming (McDougle and Taylor, 2019), RTs increased from the first rotated trial to the next trial in the 2x condition (by 42.0 ± 72.0 ms; median ± IQR; t(26)=2.94, p=0.025, d=0.55; Holm correction), but not the 1x condition (0.8 ± 53.7 ms; t(26)=0.14, p=1; Holm correction; BF_10_=0.22). On after-trials, movement direction showed a bias in the opposite direction of the rotation as evidence for implicit adaptation (**Fig. 3A, right**). A 2×3 rANOVA with the within-subject factors Condition (1x and 2x) and Rotation Magnitude (30°, 37.5°, 45°; on the previous trial) showed a main effect of Condition (1x: 3.66 ± 1.03°, 2x: 4.37° ± 1.07°, F(1,26)=11.51, p=0.002, η^2^=0.14), consistent with one additional rotated reach in the 2x condition, allowing further implicit adaptation. There was no effect of Rotation Magnitude (F(2,52)=1.35, p=0.27), which is in line with the observation that implicit learning saturates for large rotations (Kim *et al*., 2018), nor an interaction between both factors (F(2,52)=0.55, p=0.58).

Next, we examined PMBR across different trial types and conditions. By visual inspection, there was a prominent PMBR on the pre-trial, which decreased strongly in both conditions when a rotation was introduced (**Fig. 4**). PMBR was slightly decreased on the second rotated trial compared to the pre-trial. After the rotation was turned off, PMBR was similar to the pre-trial again in both conditions. A formal statistical analysis of PMBR (average across channels of interest, and the time-frequency window of interest) was based on a 2×2 rANOVA with the within-subject factors Condition (1x and 2x) and Trial Type (pre-trial and first rotated trial), thus, only including trials that were matched in kinematics. This revealed a significant main effect of Trial Type (F(1,26)=81.1, p<0.001, η^2^=0.63), a significant main effect of Condition (F(1,26)=16.11, p<0.001, η^2^=0.05), and, importantly, a significant interaction between both factors (F(1,26)=6.56, p=0.017, η^2^=0.01). Post-hoc tests revealed that there was no significant difference in PMBR between both conditions in the pre-trial (1x: 11.1 ± 14.9%, 2x: 9.0 ± 12.3%, t(26)=1.57, p=0.12, Holm correction, BF_10_=0.28), while there was a significant difference in the first rotated trial (1x:-2.4 ± 15.8%, 2x:-8.7 ± 13.8%, t(26)=4.74, p<0.001, Holm correction, d=0.44), replicating the key finding of our previous study (Korka *et al*., 2023).

**Fig. 4:**
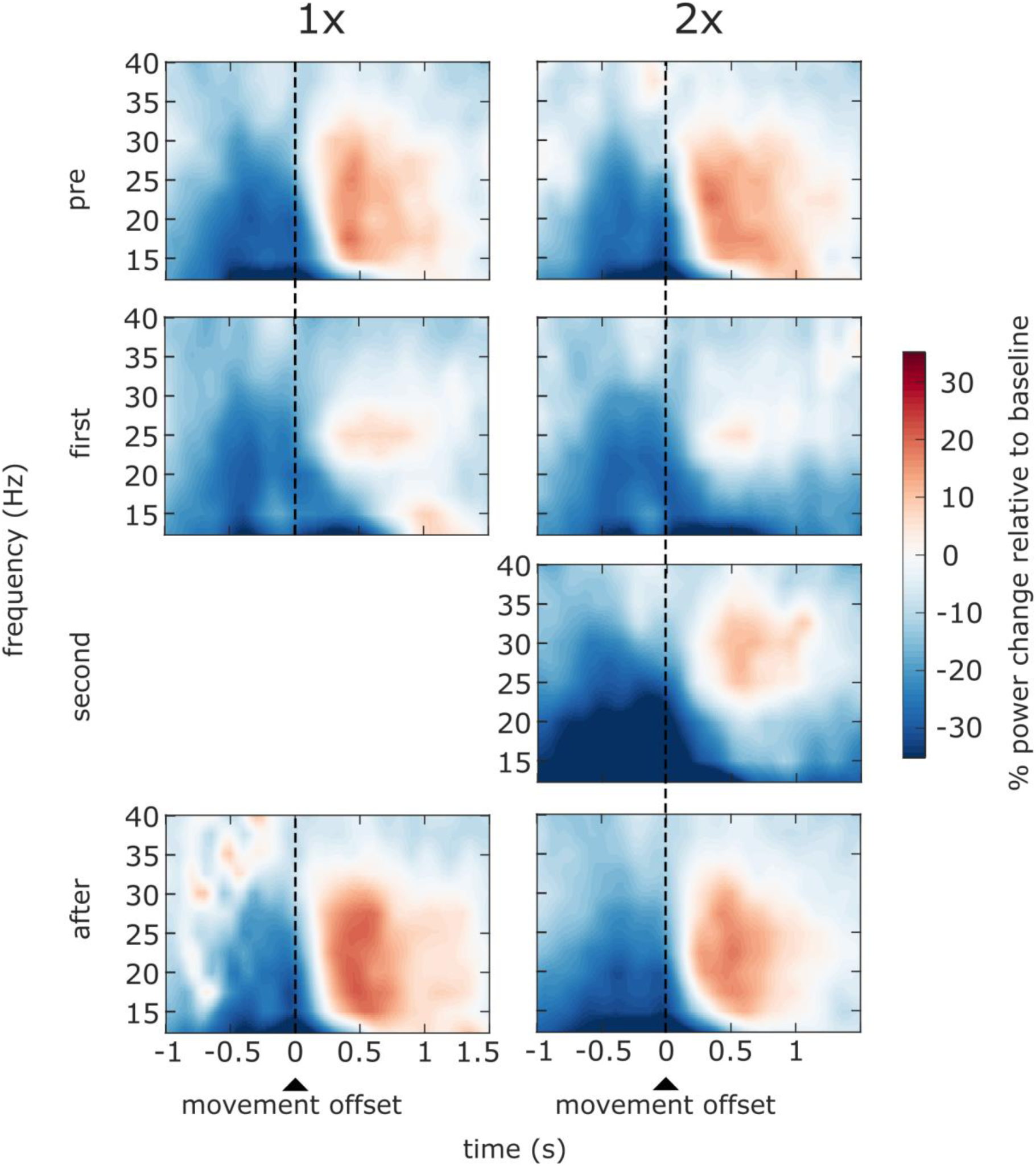
Time-frequency resolved power, averaged across channels of interest (Fig. 1A), separately for different trial types and conditions. The power in the time-frequency window of interest (200 ms to 1200 ms, 15 Hz to 30 Hz, Fig. 1B) diminished from the trial before (pre, first row) to the trial when a visuomotor rotation was first introduced (first, second row) in both conditions. The PMBR decrease was more pronounced when participants knew that the rotation would be repeated on the following trial, and they were instructed to re-aim on the second rotated trial (2x condition, right column), compared to when they knew that the rotation would not be continued and they would not have to re-aim (1x condition, left column). PMBR recovered during subsequent trials.

**Fig. 5:**
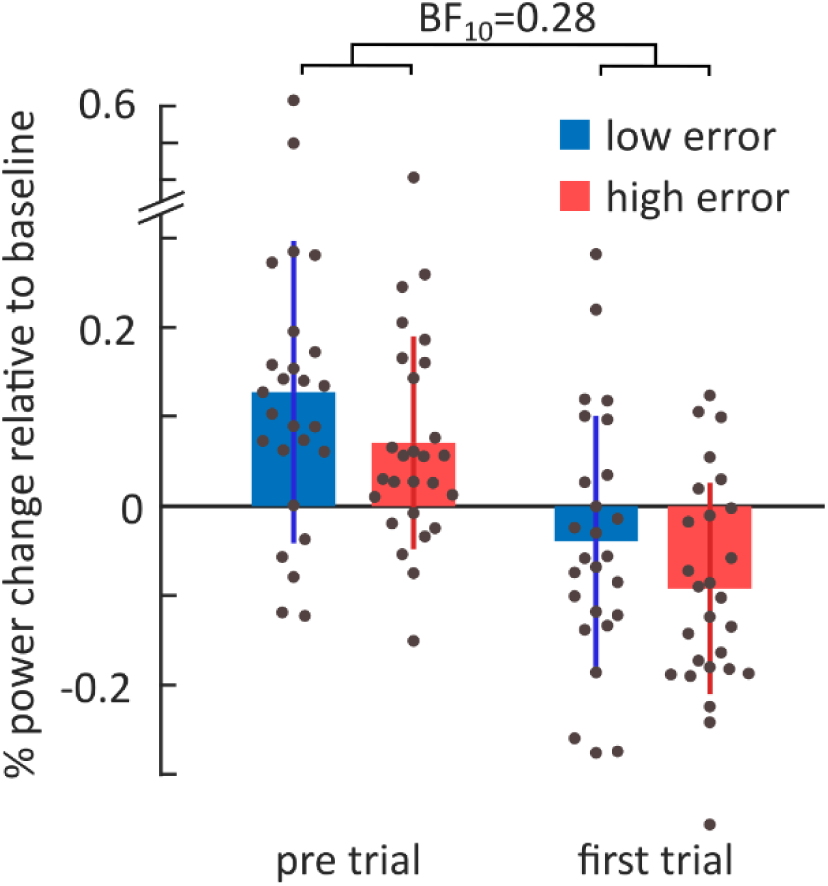
Average power change over channels and the time-frequency window of interest (bars, vertical lines represent standard deviation, each dot represents a single subject) for participants with low re-aiming error (blue) vs. high re-aiming error (red). PMBR and re-aiming accuracy were unrelated, as neither PMBR, nor its decrease from the pre-to the first rotated trial, differed between groups.

### Inter-individual differences in PMBR do not predict inter-individual differences in the accuracy of re-aiming

After confirming that PMBR was influenced by the use of an aiming strategy, we investigated whether the reduction in PMBR from the pre-trial to the first rotated trial predicts individual differences in re-aiming accuracy. To explore this, we performed a median split of participants based on their average unsigned target error in the second rotated trial, which served as an indicator of re-aiming performance. This analysis also included data from a re-aiming condition in our previous study, which was virtually identical to the 2x condition in this experiment. We first confirmed that our median-split approach was successful in separating subjects with high vs. low re-aiming accuracy. Unsigned target error on the second rotated trial was 7.00 ± 1.15° in the low-error half of subjects, and 12.50 ± 4.05° in the high-error half (t(51)=6.66, p<0.001, d=1.8). Response time costs, i.e. the increase in response time from the first to the second rotated trial, were similar across splits (low error: 13 ± 63 ms, high error: 43 ± 66 ms; median ± IQR; W=251.0, p=0.08, BF_10_=0.66), indicating that differences in unsigned error were not caused by general inattention or systematic differences in strategy approach, e.g. caching strategies versus ad hoc preparation (McDougle and Taylor, 2019).

A 2×2 mixed ANOVA with between-subject factor Re-aiming Accuracy (low error and high error) and within-subject factor Trial Type (pre-and first rotated trial) showed a main effect of Trial Type on PMBR amplitude (F(1,51)=110.5, p<0.001, η^2^=0.26), but, importantly, no effect of Re-aiming Accuracy (F(1,51)=2.5, p=0.12), nor an interaction of both factors (F(1,51)=0.02, p=0.88). Post-hoc tests also showed no significant differences in PMBR amplitude across participants subgroups (low-error vs. high-error subgroup) for the pre-trial (t(51)=1.42, p=0.16, BF_10_=0.64), first rotated trial (t(51)=1.47, p=0.15, BF_10_=0.67), or the decrease in PMBR amplitude from pre-trial to first rotated trial (t(51)=0.16, p=0.88, BF_10_=0.28).

## Discussion

We have recently shown that the decrease in PMBR during motor adaptation is amplified when participants are instructed to compensate for movement errors by using re-aiming strategies (Korka *et al*., 2023). However, it has remained unclear to what extent differences in PMBR between participants can explain differences in their ability to accurately compensate for systematic movement errors. Understanding this relationship is crucial for elucidating the role of reduced PMBR, e.g. observed in Parkinson’s disease (Pfurtscheller *et al*., 1998; Tamás *et al*., 2003) or cerebellar degeneration (Aoh *et al*., 2019; Visani *et al*., 2020), for deficits in motor adaptation. In patients with PD, reduced motor adaptation (Contreras-Vidal and Buch, 2003; Singh *et al*., 2019) is likely linked to impairments in strategic re-aiming (Tsay, Najafi, *et al*., 2022). Moreover, the cerebral cortex, basal ganglia, and cerebellum form a highly interconnected network (Hintzen, Pelzer and Tittgemeyer, 2018; Milardi *et al*., 2019). Neurodegeneration in spinocerebellar ataxia may also affect the basal ganglia, which play a key role in generating beta oscillations (Chikermane *et al*., 2024), potentially manifesting as reduced PMBR. Notably, patients with cerebellar degeneration also exhibit impairments in strategic re-aiming (Butcher *et al*., 2017; Wong *et al*., 2019). Here, we replicate the attenuating effect of strategy-use on PMBR in a paradigm that avoids potential confounds introduced by different types of perturbation in our previous study. This replication also allowed us to collapse data from the present and previous study to examine the relation between inter-individual differences in PMBR and re-aiming accuracy in a relatively large cohort of healthy subjects. Surprisingly, while PMBR shows a clear association with re-aiming at the group-level, it does not predict individual re-aiming accuracy. We discuss potential reasons for this discrepancy, together with potential future directions.

### What mental process does the modulation of the PMBR decrease relate to?

We found that PMBR decreased in response to a visuomotor rotation, an observation shared by multiple studies in the context of motor adaptation (Torrecillos *et al*., 2015; Tan, Wade and Brown, 2016; Korka *et al*., 2023). However, there is no consensus to what aspect of motor adaptation this decrease may relate to. Motor adaptation results from several interacting learning mechanisms, including implicit adaptation and strategy-based learning. The implicit adaptation system is most sensitive to presentation of visual feedback around 160 msec after movement onset, possibly corresponding to the time point of sensory prediction error computation (Wang *et al*., 2024). Here, movement duration was 310 msec on average. Thus, we would expect that neural processing related to implicit adaptation starts before movement offset. The PMBR, however, is a signal that follows movement offset, when the peak sensitivity of the implicit adaptation system to error is already over. Thus, a close association between implicit adaptation and PMBR amplitude seems a priori unlikely. On the other hand, the difference between desired and actual movement outcome, which is the driving error for strategy-based adaptation, is present upon movement offset, compatible with the timing of the PMBR. In line with this argument, delayed terminal movement feedback reduces implicit adaptation but may enhance the contribution of strategy-based adaptation (Brudner *et al*., 2016).

Here, we contrasted two conditions that engaged strategy-based motor learning to a different degree. In the 1x condition, motor adaptation was restricted to implicit learning by limiting visuomotor rotation to a single movement. In the 2x condition, adaptation resulted from a combination of implicit and strategy-based learning. We observed a stronger decrease in PMBR amplitude when participants could anticipate the same visuomotor rotation in the subsequent trial, and could, therefore, re-aim. Unlike strategy-based learning, implicit learning is not sensitive to knowledge of future perturbation (Avraham, Keizman and Shmuelof, 2020). Any neural correlate strictly related to implicit learning should thus also be invariant to expectation. On the other hand, strategy-based learning is deliberate and highly flexible, and humans exploit knowledge about the current movement context to choose optimal movement strategies (Bond and Taylor, 2015; Morehead *et al*., 2015).

Our design did not fully isolate strategy-based learning and implicit adaptation, because the latter contributed to learning in both conditions. Strategy-based adaptation interacts with implicit adaptation (Miyamoto, Wang and Smith, 2020; for review see Therrien and Wong, 2022), and the extent of implicit adaptation may, therefore, not have been exactly identical in both conditions. However, the observed pattern of PMBR differences across trial types and conditions speaks against an association between any varying degrees of implicit adaptation in the two conditions, and PMBR. Strategy-based learning typically attenuates implicit adaptation (Albert *et al*., 2022; Tsay, Haith, *et al*., 2022). If the observed decrease in PMBR amplitude upon introduction of a visuomotor rotation was related to implicit adaptation, we would, therefore, expect a less pronounced reduction in PMBR in the 2x condition, when strategy use should diminish implicit adaptation. However, we observed the opposite, i.e., a more pronounced reduction in PMBR amplitude.

Together, our results indicate that PMBR decreases more strongly when subjects re-aim. However, we found that PMBR is not predictive of individual re-aiming accuracy. A possible reason for this could be that strategy-based adaptation may unfold in separate steps. First, target error of a given movement is registered, evaluated, and stored in working memory. This likely happens following the end of a movement, when error information is available that can guide future action. Depending on the movement context, this guidance may happen at a later stage, when error information is recalled (Hillman *et al*., 2024) and helps reduce future target error (Haith, Huberdeau and Krakauer, 2015). There may be, therefore, a time gap between first processing of an error, and recall of that error for imminent action. Indeed, in our experiment, the next movement towards the target was only initiated after returning to the starting position. Thus, PMBR may not reflect action selection for the next movement, which was the return movement to the starting position, but rather processing of movement errors. Re-aiming accuracy, on the other hand, reflects both stages of strategy specification – the evaluation of error, and the specification of imminent movement. It may, therefore, not be surprising that an EEG signal that follows past movement, rather than preceding imminent movement, does not predict, by itself, behavioral performance in strategic re-aiming.

This interpretation is in line with the view that PMBR is signaling the salience of movement errors (Torrecillos *et al*., 2015). Importantly, unlike in Torrecillos et al., salience in our task was defined by the task goal, i.e., by a priori knowledge about task-relevance, rather than merely by deviance in sensory stimulation, which was constant across the 1x condition and 2x condition. We show that the PMBR decrease was amplified if participants used a movement strategy on the next trial. Importantly, movement kinematics and visual feedback of the movement were matched across conditions, at least in the critical trials (pre-trial and first rotation). Thus, any differences in PMBR between conditions necessarily relate to the movement context and its relevance for future behavior. While the target error on the first rotated trial was relevant for the task, i.e. changing movement direction on the second rotated trial of the 2x condition, it was task-irrelevant on the next (unrotated) trial of the 1x condition, as the movement direction did not change. We conclude that PMBR may be modulated by the task-relevance of errors.

Our interpretation that the modulation of PMBR codes for task-relevance of error falls well in the status quo-framework for cortical beta-band activity, which posits that beta activity is “related to the maintenance of the current sensorimotor or cognitive state” (Engel and Fries, 2010). One possible interpretation from this perspective could be that task-relevant errors signal a change in the visuomotor contingency and thus a need for adjustment of behavior for subsequent movements. Since PMBR is a physiological signal, this mechanistic explanation may be too narrowly focused on the typical structure of motor adaptation experiments. We aim to present an alternative explanation that highlights the potential ecological relevance of PMBR. In the status-quo framework, beta activity is attributed to an inhibitory role in the motor system. This aligns with findings that beta desynchronization during movement enhances cortico-spinal excitability, whereas PMBR reduces cortico-spinal excitability (Chen *et al*., 1998; Rhodes *et al*., 2024). Moreover, enhancing beta activity through transcranial alternating current stimulation has been shown to decrease movement speed (Pogosyan *et al*., 2009). Finally, successful stopping of initiated movements in a Go/NoGo task is associated with an increase in beta activity (Swann *et al*., 2009). Therefore, we speculate that a reduction in PMBR following the detection of task-relevant movement errors may alleviate motor system inhibition, enabling immediate corrective movements and thus contributing to task success.

### Limitations

Re-aiming strategies can be developed by mentally rotating the aim point for a movement in the opposite direction of the target error by an appropriate angle. Depending on working memory capacity, established strategies can be re-instantiated by caching successful responses from memory (McDougle and Taylor, 2019). We aimed to promote mental rotation and minimize response caching by increasing memory load, i.e. use of three different visuomotor rotation magnitudes in two directions, respectively. Considering the number of repetitions and use of a single target, we cannot exclude the possibility that participants started caching responses over the course of the experiment. However, this does not influence our interpretation of PMBR modulation by task-relevance of movement error as both processes should rely on the same error signal.

Is it possible that the modulation of PMBR did not result from re-aiming itself but from deviating from the usual movement direction? Movement selection should not be reflected in PMBR as it precedes movement (Alexander and Crutcher, 1990; Cisek and Kalaska, 2005).

While movement direction can be inferred from M/EEG activity during movement, direction-relevant information is encoded in low frequency activity and not in the beta band (Waldert *et al*., 2008). Indeed, beta activity is invariant across various aspects of movement (Kilavik *et al*., 2013), and thus likely not affected by movement direction.

One aim of this study was to determine the effect of different motor adaptation mechanisms on PMBR. As mentioned above, we compare two experimental conditions that engage implicit adaptation alone or implicit and strategy-based adaptation together. Thus, the effect of strategy-based adaptation on PMBR cannot be observed in isolation. On a behavioral level, both learning mechanisms can be separated by selecting appropriate feedback presentation, e.g. using movement invariant feedback to isolate implicit adaptation (Morehead *et al*., 2016), or delayed feedback to isolate strategy-based adaptation (Brudner *et al*., 2016). However, such experimental manipulations of feedback introduce differences between conditions that, by themselves, may substantially affect neural correlates of movement.

We contrasted individuals with low and high target errors to examine the relationship between PMBR and re-aiming accuracy. Inter-individual differences reflect variance that is not under experimental control. As such, its explanation requires large datasets. By collapsing datasets across two studies, we achieved a cohort size that is larger than many previous studies in the field (Tan, Jenkinson and Brown, 2014; Torrecillos *et al*., 2015; Tan, Wade and Brown, 2016). Indeed, Bayesian statistics provided moderate evidence that the main dependent variable of our median-split analysis, the decrease in PMBR amplitude from the pre-trial to the first rotated trial, was not different between individuals with high-vs. low re-aiming accuracy. However, an even larger dataset may be needed to confirm with certainty that PMBR is not directly related to re-aiming accuracy.

## Summary

We replicated our previous finding that strategy-based motor adaptation reduces PMBR amplitude. This modulation was evident even on the trial preceding the execution of a movement strategy, suggesting it may be linked to the detection of task-relevant movement errors. However, we were unable to directly associate PMBR amplitude with inter-individual differences in the accuracy of strategic re-aiming, likely due to the complex, multi-component nature of the re-aiming process. We speculate that the observed decrease in PMBR may facilitate rapid corrections of task-relevant movement errors.

## Data availability

Kinematic raw data is available under https://doi.org/10.5281/zenodo.14671785. EEG raw data is available under https://doi.org/10.5281/zenodo.14673450 and https://doi.org/10.5281/zenodo.14674428.

## Acknoledgement

We thank Fabio Dukagjini for support with data collection.

